# High-Dimensional Encoding of Movement by Single Neurons in Basal Ganglia Output

**DOI:** 10.1101/2023.05.17.541090

**Authors:** Gil Zur, Noga Larry, Mati Joshua

**Affiliations:** Edmond and Lily Safra Center for Brain Sciences, the Hebrew University, Jerusalem, Israel

## Abstract

The Substantia Nigra pars reticulata (SNpr), an output structure of the basal ganglia, has been hypothesized to gate the execution of movements. Previous studies focusing mostly on saccadic eye movements have reported that SNpr neurons are tonically active and either pause or increase their firing during movements, consistent with the gating role. We recorded activity in the SNpr of two monkeys during smooth pursuit and saccadic eye movements. SNpr neurons exhibited highly diverse reaction patterns during pursuit, including frequent increases and decreases in firing rate, uncorrelated responses in different movement directions and in reward conditions resulting in the high dimensional activity of single neurons. These diverse temporal patterns surpassed those in other oculomotor areas in the frontal cortex, basal ganglia, and cerebellum. These results suggest that temporal properties of the responses enrich the coding capacity of the basal ganglia output beyond gating or permitting movement.

## Introduction

The basal ganglia are embedded within cortical and sub-cortical networks that drive movement^1^. Initial studies proposed that the basal ganglia directly control behavior, with tonic inhibition limiting movement and pauses in basal ganglia activity facilitating movement initiation^2, 3^. However, rather than a binary coding of movement, the output of the basal ganglia coded behavioral events by both increases and decreases in activity^4, 5^. One explanation was that this pattern of activity might reflect the pattern of inhibition and excitation (disinhibition) required to drive movement^6, 7^. Alternatively, the complexity of single neurons might not conform directly to movement parameters as complexity could emerge from internal computations, such as those that are found in the implementation of a dynamical system^8, 9^. Complex temporal patterns of responses of single neurons that are not directly related to movement parameters are a signature of such dynamics. Therefore, a critical question in basal ganglia research is whether the temporal patterns of activity are complex, and how they related to movement parameters.

In the current study we approached this issue by exploring the temporal pattern of the activity of single neurons in the basal ganglia during behavior. We focused on the eye movement system since it provides exquisite control over behavior, as shown by decades of research that have characterized responses in the final motor pathways^10–12^ thereby providing a framework for interpreting the observed temporal activity patterns. Previous research has mostly studied basal ganglia activity during saccadic eye movements^13–15^. However, the rapidity of saccades constrains the ability to identify temporal modulations in movement coding. In contrast, smooth pursuit eye movements are continuous and prolonged, which provides an opportunity to examine modulations that may be indiscernible during saccades. To examine the potential temporal modulations in movement coding, we perform recordings from the basal ganglia during both saccadic and pursuit eye movements.

We recorded activity in the substantia nigra pars reticulata (SNpr), the basal ganglia output structure that is involved in both pursuit and saccade eye movements^16, 17^. We show that single neural activity exhibits diverse temporal response patterns. These manifested in the weak correlations between responses and the high dimensionality of single-neuron activity across the direction of pursuit eye movements. Notably, the diversity of activity patterns was greater in the SNpr compared to other oculomotor brain areas, including the caudate, cerebellum, and frontal eye field. The findings emphasize the temporal characteristics of SNpr neurons, underscoring the importance of temporal coding in elucidating the function of basal ganglia output. The contrast between the complex temporal patterns and simple pursuit behavior suggests that rather than coding movement parameters through inhibition or disinhibition, the basal ganglia extracts information that resembles intermediate stages of neural networks.

## Results

### Behavioral task and neural recordings

Two monkeys were trained on saccade and pursuit tasks in which we manipulated the reward probability (Figure 1A). Both tasks were initiated by a fixation stage in which a white fixation target appeared in the center of a black screen. After 500 ms, the target changed to one of two colors, indicating the probability of the reward given after the successful completion of the trial. One color corresponded to a probability of 75% and the other to a probability of 25%. After a random period (800-1200 ms) the target moved in one of eight directions. In the saccade task, the target instantaneously jumped to a 10° eccentric position. In the pursuit task, the target moved continuously at a speed of 20°/s^18^. The monkeys were required to track the target accurately to complete the trial. At the end of a successful trial, the monkeys received the reward corresponding to the probability specified by the color of the target. Figure 1B presents examples of horizontal eye positions on trials where the target moved to the right. In the saccade task, the monkeys saccaded rapidly to the new location of the target, whereas on the pursuit task the monkeys followed the target smoothly until it stopped.

**Figure 1:**
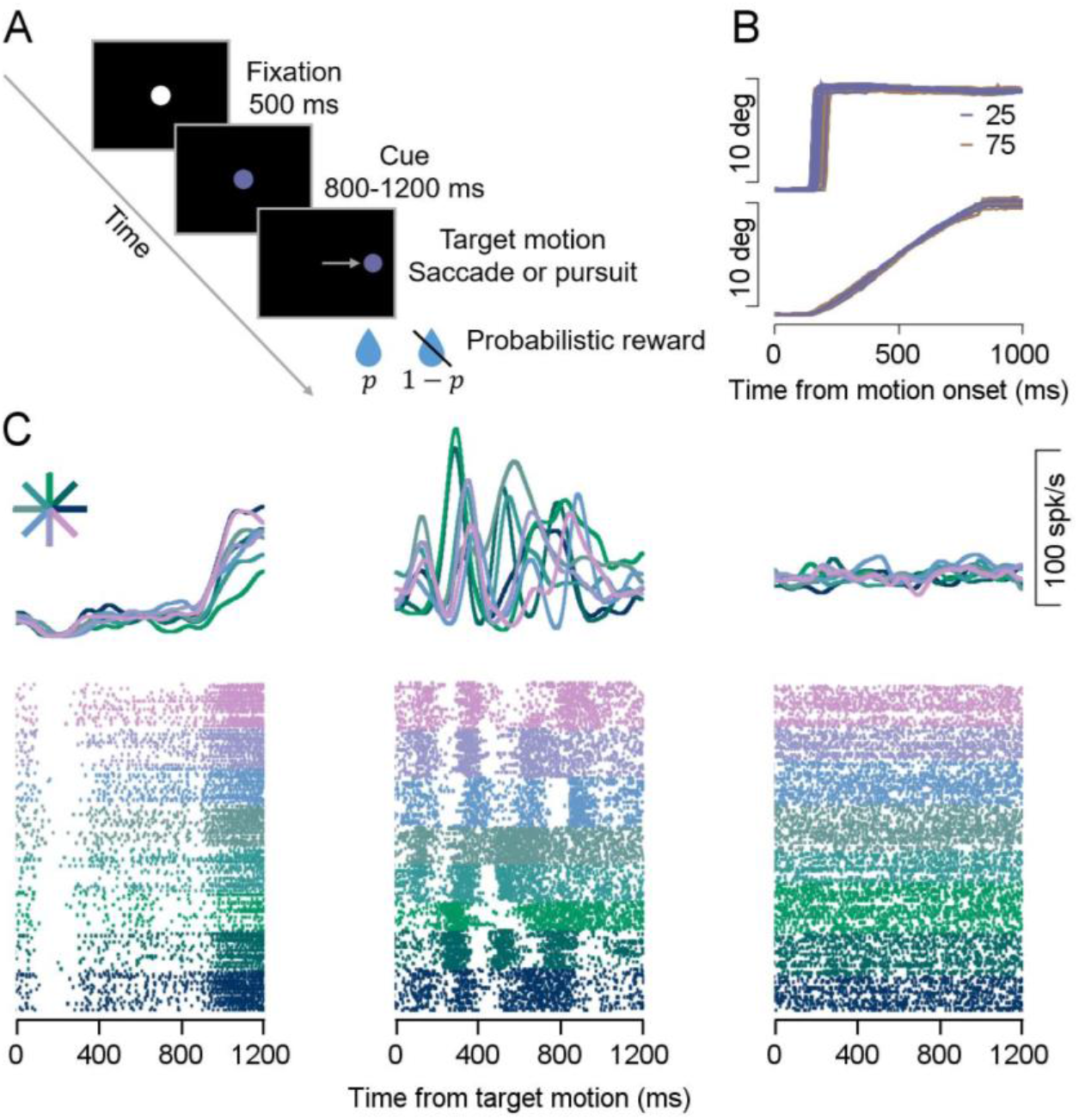
Behavioral task and example neurons. **A,** Schematics of the task stages as they appear on the screen: fixation target, color switch to cue reward probability, and target motion in one out of eight directions. A reward was delivered after completing the trial according to the probability (p) associated with the cue. **B,** Horizontal eye position for several saccades (top) and pursuit (bottom) trials in which the target moved to the right. The horizontal and vertical axis represents the time from motion onset and the eye position in degrees. The colors indicate the probability to receive the reward. **C,** PSTH (top), and raster (bottom) of three example neurons. Each column shows a single neuron during target motion in a pursuit session. Colors indicate the direction of the target motion and correspond to the direction presented in the asterisk at the top of **C**.

We recorded from the monkeys’ SNpr while they performed the eye movement tasks. Figure 1C shows the averages and trial-by-trial activity of example SNpr neurons recorded during the pursuit task. Since the effect of reward on behavior and neural activity during movement (but not the cue) was very small^19^, we initially grouped trials in the two reward probability conditions based on the direction of movement. Later we will address the effects of the reward probability in detail. The first neuron consistently paused its activity after the initiation of movement in all directions and trials (Figure 1C left). This pattern of activity is similar to findings elsewhere indicating that tonically active SNpr neurons exhibit either a pause or an increase during eye movements ^20–22^. The second type of activity was characterized by a diverse reaction pattern across directions of movement (Figure 1C middle). Importantly, examination of the raster indicated that these multiphasic increases and decreases in firing rate were consistent across trials having the same direction of movement. By contrast, a neuron that did not respond to the task (Figure 1C right), was characterized by a inconsistent modulation between different trials, resulting in low average amplitude activity in the PSTH. Thus, these examples demonstrate that in our dataset some neurons responded stereotypically as reported previously whereas others exhibited diverse temporal patterns across directions of movement.

### Characterizing the diversity of temporal patterns

To quantify the temporal diversity of responses in the SNpr, we calculated the correlation between activity on different trials (see Methods for more details). Figure 2 depicts a neuron presenting a diverse temporal pattern (Figure 2A). We first smoothed the activity in each trial with a Gaussian filter (SD 30 ms, see Figure S1 for other SDs) and then calculated the correlation matrix between all trials (Figure 2B). Trials were ordered by direction of movement. Hence the pattern of squares along the diagonal in the correlation matrix indicates that trials from the same condition tended to have a correlated response pattern. The squares off-diagonal show the correlations under different conditions. For each two conditions, we defined the *Between* score as the average of all the single-trial correlations between conditions. This is depicted in Figure 2C in the black square representing the mask of all the values averaged for conditions *i* and *j.* We compared the *Between* score to the *Within* score, which was defined as the geometric mean of the average correlation within each condition (excluding the correlation of a trial with itself). This is represented in Figure 2C by the triangles near the diagonal that mask the values averaged to calculate the *Within* score. In the Appendix, we show that the ratio between the *Between* and *Within* scores equals the correlation between the average temporal pattern of the responses. Therefore, a ratio of 1 indicates linear scaling of the responses across conditions whereas larger *Within* than *Between* scores characterize diverse temporal patterns across conditions.

**Figure 2:**
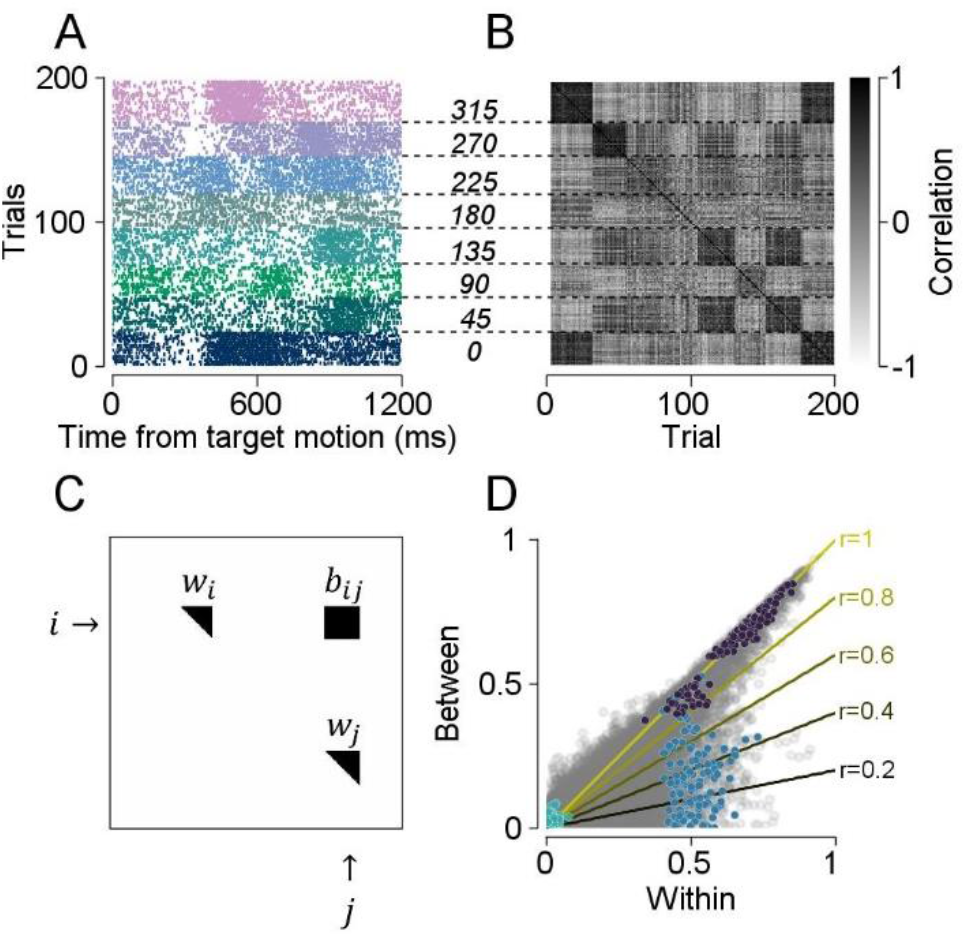
Diversity of SNpr responses as indicated by the Within vs. Between scores. **A,** Raster of a single neuron. Each row is the activity of a single trial during the target motion. Dashed lines from **A** to **B** mark the condition borders with the direction of target movement in degrees. Colors represent the different directions of movement. **B,** Correlation matrix calculated over the rows of the raster matrix in **A** after the raster was smoothed with a Gaussian filter. The brightness of each pixel indicates the size of the correlation. **C,** Mask over the correlation matrix in **B**. *i* and *j* are two different conditions. The black square (labeled *b_ij_*) marks the position in the correlation matrix shown in **B** that corresponds to the correlation of trials *between* the *i*th and *j*th condition. The triangles mark the position in the correlation matrix of the correlation *within* the *i*th and *j*th condition (labeled as *w_i_* and *w_j_*). **D,** Scatter of Within (horizontal axis) versus Between scores (vertical axis) for all neurons from the SNpr. Each dot represents a pair of conditions from individual neurons from the SNpr. Colored dots are example neurons from Figure 1C. Dark blue, blue, and green correspond to the neurons on the left, middle, and right of Figure 1C. Solid lines indicate different theoretical correlation slopes specified by the r value (see Appendix).

Figure 2D shows the *Within* versus *Between* scores for all neurons in the SNpr and all pairs of direction conditions. Responses consistent (exhibiting linear scaling) across conditions resulted in large *Within* and *Between* scores. These fall near the equality line, far from the origin (Figure 2D dark dots showing neuron 1 from Figure 1C). Responses with diverse temporal patterns across conditions resulted in large *Within* and small *Between* scores. These appear beneath the equality line, far from the origin (2D in blue showing neuron 2 from Figure 1C). Finally, neurons with small responses had small *Within* and *Between* scores. These fell near the origin (2D in green showing neuron 3 from Figure 1C). Thus, the comparison of the *Within* and *Between* scores serves to differentiate the three patterns shown in Figure 1. Neurons with responses that scale linearly plot along the diagonal, neurons with diverse temporal patterns plot beneath the diagonal, and neurons with small responses plot near the origin. Note that the latter two cannot be distinguished from the calculation of the correlations of the average activity.

Overall, we found numerous responses from the SNpr that fell far from the origin beneath the equality line, indicating that neurons in the SNpr encoded different directions of movement with diverse temporal patterns (Figure 2D). We compared the pattern of scores to the pattern expected from different values of the correlation of average activity (lines in Figure 2D). We found that many pairs of conditions matched the low average correlations. This demonstrates that many SNpr neurons coded movement direction with a rich temporal pattern of activity, diverging from the simple characteristic increase/decrease in activity that is classically assumed to gate or enable movement.

### SNpr responses are more diverse than other cortical and subcortical populations

To test whether the diverse responses in the SNpr are specific or whether other cortical and subcortical populations also have similarly diverse responses, we calculated the *Within* and *Between* scores of neurons in other oculomotor areas recorded in the same monkeys: the input structure of the basal ganglia (striatum) and the vermis of the cerebellum^19^. We also compared these results to neurons from the floccular complex of the cerebellum^23^ and frontal eye field (FEF) recorded in similar tasks in other monkeys^6^.

We binned the *Within* score (i.e., Figure 2D) into quantiles and then computed the mean *Within* minus *Between* differences for each quantile. We denote this difference between scores (*Within-Between*) as the *Diversity score* since larger values indicate more diverse responses across conditions. Figure 3A shows the relationship between the *Diversity* score and the *Within* score during pursuit (see Figure S2A for the saccade task). This presentation highlights the regime of strong responses (*Within* >> 0) in which there was a clear difference in the diversity scores. We found that the diversity of the SNpr during pursuit was significantly higher than in other regions in the eye movement pathway (Figure 3A, SNpr vs. Striatum p=7.9e-37, SNpr vs. Vermis p=1.3e-92, SNpr vs. Flocculus p=3.8e-46, SNpr vs. FEF p=4.4e-47 Wilcoxon signed-rank test, see Methods). We also found that for very large *Within* scores, the temporal diversity of SNpr decreased. This indicates that in addition to the complex responses in the SNpr there is a subpopulation with very large and stereotypic responses (e.g., Figure 2D, dark dots).

**Figure 3:**
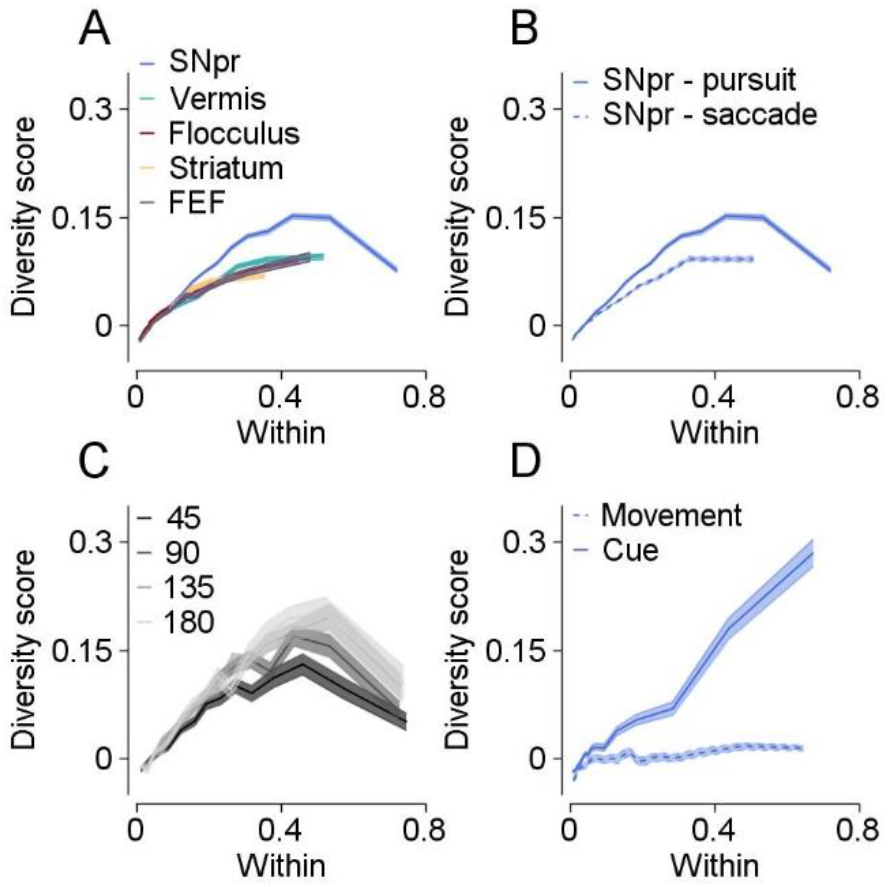
Temporal diversity of SNpr neurons exceeds other populations. **A-D,** Each plot shows the Diversity scores (Within-Between, vertical axis), as a function of the Within score (horizontal axis). The horizontal axis was binned into quantiles (15 bins). **A.** Comparison between regions. Colors indicate different neural populations. **B,** Comparison between saccade (dashed line) and pursuit (solid line) tasks in the SNpr. **C,** Comparison between distances in target motion direction. Colors indicate the angle between conditions. **D,** Comparison of reward probability diversity during cue (dashed line) and movement (solid line).

### The diversity of responses depends on sensorimotor parameters

The SNpr responses were more diverse during pursuit than during saccades (Figure 3B, p=1.8e-58, Wilcoxon signed-rank test see Methods), indicating that diversity depends on the sensorimotor profile of the behavior (but not on the pattern of catchup saccade during pursuit, Figure. S2B). To further characterize the relationship between diversity and the sensorimotor properties of the movement we compared similarity in temporal pattern as a function of similarity in movement direction. The diversity of the response increased with the difference in the angle of movement (Figure 3C, see Figure S2C for saccade).

Finally, we also compared the response activity across different reward conditions. We presented two conditions of reward probability (25% and 75% of receiving a reward). Data were collected from both conditions in all eight directions of movement. As expected from the small reward-probability modulations during movement^19^ we did not find diverse reward-related responses, as indicated by the low diversity scores across reward probability conditions (Figure 3D, dashed). This also confirms the validity of our decision to group responses with different reward probabilities. By contrast, at the cue epoch when the color target appeared and the monkeys were fixating on the target, we found diverse responses (Figure 3D, solid, see Figure S3 for other populations) thus indicating that diversity in patterns of activity is not limited to the coding of movement direction.

### Multidimensional responses of single neurons in the SNpr

Low correlation in the pairs of neural responses suggests that the neural responses across movement directions cannot be described by a single temporal pattern. Thus, the neural code for movement direction is likely to contain a large number of distinct temporal patterns of activity. These temporal patterns can be conceptualized as dimensions within the activity space of a single neuron, whereby a higher dimensionality signifies an increased capacity for representing a greater range of functions within the activity space. To quantify this dimensionality, we developed an analysis to estimate the number of independent dimensions required to explain the temporal pattern of the neuronal response (Figure 4E, see Methods). This analysis involved first computing the PSTH of each condition. We then calculated the total variance in the PSTH matrix (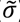) to serve as a statistic for computing the dimensionality. To assess the significance of 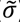, we generated a distribution of variances by shuffling the trials’ conditions before computing the PSTHs. If 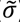 was significant, the subsequent step involved the progression to the next dimension by eliminating activity from the neural responses in the direction exhibiting the highest variance (1^st^ principal component) within the PSTH matrix. This process continued until the variance across PSTHs was no longer significant, yielding an estimate of the number of dimensions required to explain the variability of the neuronal response. We validated through simulations that across a broad spectrum of noise conditions the algorithm identifies the correct number of dimensions (Figure S4). In this analysis, stereotypic responses in all directions have a dimension of 0. Any responses characterized by linear scaling (such as the characteristic examples of tuning curves^24^) have a dimension of 1.

**Figure 4:**
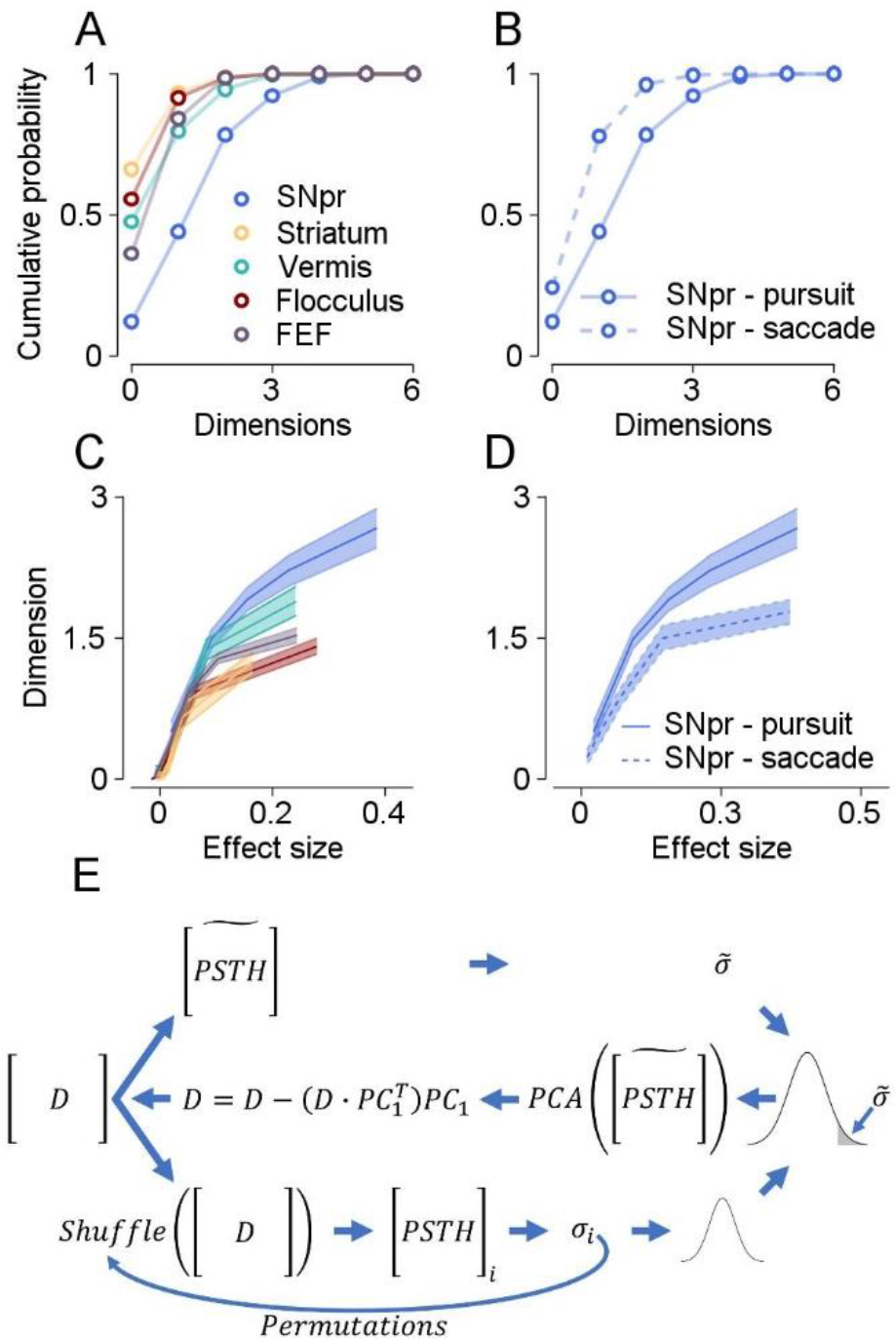
Dimensionality of neurons in the SNpr exceeds other populations. **A.** Cumulative distribution of the dimension of the neurons. Colors indicate different populations. **B**. Cumulative distribution of the dimension in the SNpr during the saccade (dashed line) and pursuit (solid) tasks. **C-D,** Dimension of neurons (vertical axis) as a function of effect size (horizontal axis). Colors same as in **A** and **B**. **C,** Comparison between different populations. **D,** Comparison between saccade (dashed line) and pursuit (solid line). The horizontal axis was binned into quantiles (15 bins). **E.** Dimension analysis illustrated in a diagram, where arrows indicate the sequential steps of the analysis. D matrix represents the data matrix of all trials across time. The upper part shows the computation of the PSTH and statistic marked with tilde. Lower part represents the computation of a distribution for the statistical test. The *i*th PSTH and sigma are generated by shuffling the D conditions, and a distribution of sigma is created. The sigma tilde is compared to the distribution (middle row on the right), and if significant, the first PC (PC1) of PSTH tilde is extracted and all the variance of D in the direction of PC1 is subtracted (middle). The analysis is repeated with the updated D matrix.

We found that 56% of the SNpr neurons had a dimension of 2 or more during pursuit (Figure 4A, see Figure S5 for saccade). The SNpr trace plotted the lowest in the cumulative distribution of dimensions, indicating that the activity of SNpr neurons had higher dimensions than the other populations (Figure 4A, SNpr vs. vermis p=8.4e-15, SNpr vs. flocculus p=2.2e-27, SNpr vs. MSN p=2.0e-26, SNpr vs. FEF p=1.2e-18, Wilcoxon rank-sum), enabling them to span a broader range of functions. Moreover, the dimension analysis for the SNpr neurons revealed a higher dimension during the pursuit than during saccades (Figure 4B, SNpr pursuit vs. SNpr saccade p=2.3e-7, Wilcoxon rank-sum), indicating that individual SNpr neuron diversity depends on the sensorimotor profile of behavior and not merely on time.

To confirm that the higher dimensions observed in the SNpr were not simply a result of larger responses, we calculated the effect size (see Methods)^19^ of each neuron and compared the dimensions of the SNpr to those of other brain areas involved in eye movements. We found that even when controlling for effect size, SNpr neurons had significantly higher dimensions than the other populations (Figure 4C, SNpr vs. MSN p=2.6e-5, SNpr vs. FEF p=2.1e-8, SNpr vs. vermis p=0.01, SNpr vs. flocculus p=7.1e-8, Wilcoxon signed-rank test, see Methods). Comparing the dimensions of the SNpr neurons for similar effect sizes during the pursuit and saccade tasks yielded a higher dimensionality in the responses when the monkeys performed pursuit eye movement task (Figure 4D, p=9.1e-7 Wilcoxon signed-rank test, see Methods). Thus, SNpr temporal diversity is related to the behavior, with higher diversity during pursuit. During saccades the difference between populations was less pronounced (Figure S5), indicating that the examination of pursuit behavior was key to revealing the access diversity of the SNpr.

## Discussion

We found that the SNpr neurons exhibited highly diverse reaction patterns during movement, including frequent increases and decreases in firing rate, as well as uncorrelated responses in different directions and reward conditions resulting in the high dimensional activity of single neurons. The activity in SNpr was more diverse than in other motor areas. Furthermore, the pattern of SNpr activity differed from patterns of eye motoneuronal activity which are linearly related to the projection of the eye velocity and position in the preferred direction of muscle activity (Figure S6)^25–27^. Thus, the activity in the SNpr does not match the pattern needed to excite or inhibit single motoneuronal activity. Instead, our results suggest a dynamic complex mapping between sensorimotor parameters and activity, akin to intermediate levels in artificial neural networks. Diverse and high dimensional responses in neurons are beneficial, since they provide the possibility for extensive classification through simple linear readouts^28,29,30^, and can encompass a wide range of behavioral functions. Our results suggest that the SNpr is capable of engaging in the separation and classification of task-related information, and embedding it in a high-dimensional space that creates opportunities for versatile and diverse high-level task performance.

The mechanism underlying the emergence of diversity in the basal ganglia remains a topic of investigation. Studies of the arm movement system have postulated that diversity arises from an autonomous dynamical system implemented through local recurrent connectivity^31, 32^ (but see ^33^). However, unlike the arm movement system, the initiation and maintenance of the pursuit system heavily rely on visual inputs, suggesting that the observed diversity during pursuit is more likely to arise from the way visual inputs are processed and integrated by the neural circuitry rather than being solely dependent on local recurrent connectivity. Although the basal ganglia exhibit local reverberating connectivity that could generate temporal diversity, the output nuclei are mostly driven by feedforward inputs via direct and indirect pathways from the striatum and the hyper-direct pathways from the cortex^34^. Thus, our results suggest that diversity in output is achieved through connectivity between areas rather than local recurrent connectivity. The lower diversity of the striatal neuron (Figures 3A and 4C) suggests that diversity arises from the pattern of convergence. A regression analysis (not shown) further confirmed that diversity responses in the SNpr can be predicted from the many striatal neurons supporting the role of inputs rather than de novo generation of diversity in the SNpr.

The results of the current study holds promise for further exploration in several critical directions. We studied diversity in the coding of movement direction during saccades and pursuit. Future research could examine the ways in which diversity is related to other parameters of movement, such as duration or speed and probe more densely the direction of movement. We found diverse responses even in the closest direction of movement (Figure 3C), suggesting discontinuity in the coding of movement direction. Identifying and characterizing these discontinuities could provide further insight into the coding of movement direction. In addition, we compared diversity in the SNpr to inputs from the striatum and other brain regions outside the basal ganglia. A compelling avenue for future research would be to compare this diversity to outputs of the SNpr, specifically comparing with the superior colliculus and thalamus to study how the diverse activity is readout by downstream structures. Finally, further characterization of the inputs to the SNpr such as dissection of the inputs from the external segment of the globus pallidus or subthalamic nucleus or selective probing of direct and indirect pathways^35, 36^ could characterize how diversity arises within the basal ganglia.

## Methods

All experimental procedures were approved by the Institutional Animal Care and Use Committees of the Hebrew University of Jerusalem. Data were collected from one female (Monkey G) and one male (Monkey A) Macaca Fascicularis monkeys that had been prepared for behavior, and neural recording using techniques described in details previously^19^. Briefly, we implanted head holders to restrain the monkeys’ heads and trained the monkeys to track spots of light that moved across a video monitor positioned in front of them (55 cm and 63 cm from the eyes of monkeys A and G). We used liquid food rewards (baby food mixed with water and infant formula) from a tube placed in front of them to reward monkeys for accurate tracking of the targets.

The position of the eye was continuously tracked using a high temporal resolution camera (Eye link - SR research) at a frequency of 1 kHz and the data were collected for further analysis. To record from the basal ganglia and the vermis, we performed two subsequent surgeries to place a 19 mm diameter cylindrical recording chamber over these brain regions. We first recorded from the caudate and then moved on to the SNpr, while continuously recording from the vermis. Based on an extensive mapping of the caudate and an MRI image of the brain, we estimated the location of the SNpr in the recording chamber. Then, we lowered the electrodes to this location. At the targeted site, we identified neurons with a high baseline firing rate and the typical extracellular shape of SNpr neurons^37^. We confirmed that some of these neurons exhibited a clear pause during saccades in certain directions. On some recording days, we also identified neurons with pronounced eye position sensitivity, as expected from the neighboring third oculomotor nerve. We used two Mini- Matrix Systems (Thomas Recording GmbH) to lower quartz-insulated tungsten electrodes (impedance of 1–2 Mohm) into the basal ganglia and cerebellum. The neural signals were digitized at a sampling rate of 40 kHz using the OmniPlex system (Plexon) and sorted offline (Plexon spike sorter). We only used neurons that displayed distinct clusters of waveforms in the sorting procedure. The sorted spikes were converted into time stamps with a resolution of 1 ms and were inspected again visually to identify any instability or obvious errors in the sorting procedure. To control for potential behavioral differences between reward conditions that might affect our results, we also recorded the monkeys’ licking behavior. We used an infrared beam to track the licks. Monkey A tended not to extend its tongue, so we recorded its lip movements instead.

### Experimental design

Before beginning the neural recordings, we examined the behavior of the monkeys under different reward probability conditions. On another task in which the monkeys were required to choose between two targets, in more than 97% of the trials the monkeys chose the 0.75 over the 0.25 probability target, indicating that they associated the color of the target with the probability of reward^19^.

### Pursuit Task

Each trial started with a bright white target that appeared in the center of the screen (Figure 1A). After 500 ms of presentation, in which the monkey was required to acquire and maintain fixation, a colored target replaced the fixation target. The color of the target signaled the probability of receiving a reward upon successful tracking of the target. For monkey A we used yellow to indicate a 75% for reward and green to indicate 25%. For monkey G we reversed the associations. After a variable delay of 800-1200 ms, the targets stepped in one of eight directions (0°, 45°, 90°, 135°, 180°, 225°, 270°, 315°; the same directions were used for all tasks below) and then moved continuously in the opposite direction (step-ramp)^38^. For both monkeys, we used a target motion of 20°/s and a step to a position 4° from the center of the screen. The target moved for 750 ms for monkey A and 650 ms for monkey G and then stopped and stayed still for an additional 500-700 ms. If the monkey’s gaze was within a 3-5°x3-5° window around the target, the monkey received a reward according to the probability specified by the color.

### Saccade Task

The structure of the saccade task was identical to the pursuit task, except for the target motion epoch. In the saccade task, following the random delay, the central colored target disappeared and immediately reappeared in one of eight eccentric locations 10° from the center of the screen. If the monkey’s gaze was within a 5°x5° window around the target, the monkey received a reward according to the probability specified by the color.

### Flocculus reward size task

We used previously analyzed data recorded from the floccular complex and adjacent areas, while the monkeys performed a smooth pursuit task in which we manipulated reward size. The full details can be found in previous reports^39, 40^. Briefly, the temporal and target motion properties of the task were the same as the reward probability pursuit task described above. However, in this task, the color of the target signaled the size of the reward the monkey would receive if it tracked the target. One color was associated with a large reward (∼0.2 ml) and the other with a small reward (∼0.05 ml).

### FEF task

We analyzed two types of trials recorded from one monkey in the FEF. In the pursuit task, the trials began with a fixation target displayed in the center of the screen. After 500 milliseconds, an eccentric cue appeared at one of eight locations four degrees away from the fixation target. This eccentric cue indicated the opposite direction of the future movement of the target. After a random delay ranging from 800 to 1200 milliseconds, the target began to move in the direction indicated by the eccentric cue at a speed of 20 degrees per second for 750 ms. In the saccade task, the trials began with a fixation target displayed on the screen for a random period of time ranging from 1300 to 1700 milliseconds. Then the target jumped to one of eight eccentric locations 10° away from the center of the screen. In both tasks, the monkey received a reward at the end of the trial if it successfully followed the target.

### Data Analysis

Analyses were performed using Python and Matlab. To ensure accuracy, we only considered cells that had been recorded for a minimum of 100 trials in either the saccade or pursuit task. Since there were only 80 available trials in the FEF region, we conducted a control analysis comparing the FEF to the SNpr region with a similar number of trials (Figure S7).

**Table 1:**
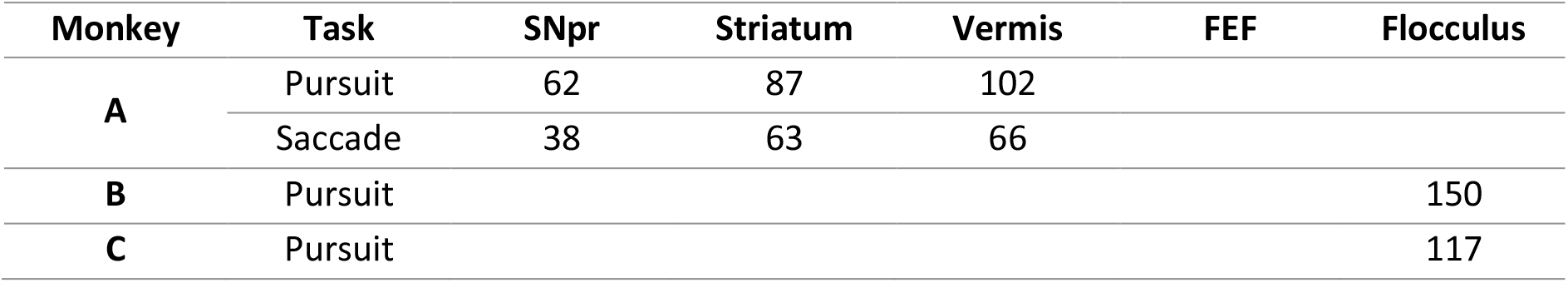
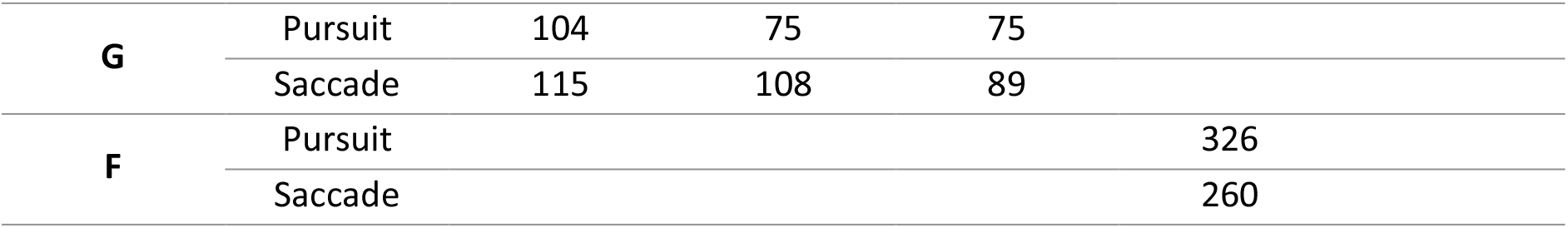
Number of neurons analyzed in each task and area of recordings

Given the dynamic nature of the analyses, we paid particular attention to timing and focused on a 1200 ms time window starting from the target movement onset. We evaluated the impact of taking shorter time windows after the end of the motion epoch (such as 200, 400 ms) to ensure that our results were not sensitive to this window, especially in the saccade task where movements are shorter.

### Saccade detection

We used eye velocity and acceleration thresholds to detect saccades automatically which was then verified by visual inspection of the traces. The velocity and acceleration signals were obtained by digitally differentiating the position signal after smoothing with a Gaussian filter with a standard deviation of 5 ms. Saccades were defined as an eye acceleration exceeding 1000°/s^2^, an eye velocity crossing 15°/s during fixation, or an eye velocity crossing 50°/s while the target moved on the pursuit task.

### PSTH calculation

To calculate the PSTH of a neuron, we first smoothed all the trials with a Gaussian filter where the standard deviation was 30 (see Figure S1 for control with other kernel sizes). We then grouped the trials by condition and calculated the average for each group for each time point.

### Analysis of Between and Within scores

We compared each pair of the task conditions for each neuron. For each neuron, we began our analysis with its *n*×*m* raster matrix, where *n* is the number of trials, and *m* is the number of time points in ms. We began by smoothing the activity in each trial with a Gaussian filter and then calculated the Pearson correlation matrix between all trials (i.e., row-wise correlation).

The correlation matrix was then used to define the *Within* and *Between* scores for each pair of conditions *i* and j as follows:

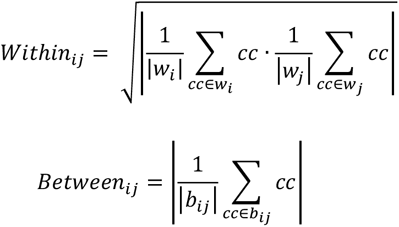

where *<u>w_i_</u>* is the group of all the correlations across trials in condition *i* (excluding correlations of a trial with itself), and *b_ij_* is the group of all correlations between trials in the *i*-th condition and trials in the *j*-th condition. We define the *Diversity* score as the *Within* minus *Between* score (*Within* -*Between*).

### Testing neuronal measures using score matching

In the results section, we showed that for a given *Within* score, the *Between* score of the SNpr neurons tended to be smaller (Figure 3). We next compared the *Between* score of two populations while controlling for the *Within* score. We first matched each *Within* score in one population to the closest *Within* score in the other population. We then applied a Wilcoxon signed-rank for paired samples test to compare the *Between* scores of the responses that were matched by the *Within* score. Statistical significance in this test indicates that the null hypothesis that responses in the two populations with the same *Within* score have the same *Between* score is unlikely. In this analysis, we matched the population with a larger range of *Within* scores to the population with a smaller range to ensure that for each *Within* score in one population we would be able to find a close *Within* score in the other population. After matching there were no differences between the *Within* scores of the matched population (p>0.15, Wilcoxon signed-rank test).

We used this test to compare the SNpr to the other populations in the *Within*-*Between* analysis (Figure 3) and to compare the SNpr during pursuit and saccade. We used a similar test to compare the dimension as a function of the effect size (Figure 4), with the effect size of the neurons with dimension replacing the *Within* and *Between* Score.

### Dimension Analysis

We defined the dimension (*d*) of a neuron as the number of significant dimensions by Principal Component Analysis (PCA) over the PSTH of that neuron, as follows: let *D* be a data matrix of a neuron, where each row of *D* is the activity of the neuron during the segment of interest from a single trial. Averaging over columns of *D* for rows with the same trial condition result in a new matrix *PSTH_real_*, where each row is the average activity in time during a single condition. The *PSTH_real_* matrix is then binned at a 100-ms interval, and the average activity in each bin is calculated.

To test for the significance of the first dimension, we first calculated the total variance across *PSTH_real_*. Next, we built a distribution of variances (1000 iterations) over multiple shuffled PSTH matrices. A shuffled PSTH matrix was generated by shuffling the trials’ conditions of *D* before averaging the PSTH as described above. The dimension was significant (i.e., d incremented by 1) if the total variance was larger than 95% of the variances generated by the shuffled PSTH matrices (0.05 significance level). We then performed principle component analysis (PCA) on the *PSTH_real_* and removed all the variances in the direction of the first PC from each trial in the original data matrix (*D*). This was done by subtracting the projection in the direction of the first principal component of each trial:

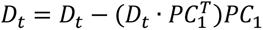

where t is the index of the trial. By subtracting, we eliminated all the variance in the first PC direction. This enabled us to repeat the above analysis on the resulting data matrix *D*, where the first eigenvector was the previous second eigenvector. We proceeded until the variance across PSTHs stopped being significant. The number of dimensions that passed the significance test (*d*) was defined as the dimension of the neuron. The whole process is illustrated in a diagram in Figure 4E.

To further validate our methods, we conducted a simulation to test the performance of response dimension method (Figure S4). We generated neurons with the same number of conditions as in our task (conditions=16) with 10 trials per condition. The signal of each neuron was generated with a number (0-6) of orthogonal dimensions using cosine and sine functions with different frequencies; i.e., cos(*N·t*) and sin(*N·t*), where *t* represents the time between 0 and 2π and *N* is an integer. We added Gaussian noise (sigma range from 0.1 to 1.5) to test for the accuracy of the analysis in the presence of noise. We calculated the effect size of the simulated neurons to ensure that the dimension analyses performed well in the regime of trial-by-trial variability in the activity of the recorded neurons. The results indicated satisfactory prediction of the dimension for effect sizes exceeding ∼0.2 (see Figure S4) and a monotonic relationship between the true and calculated dimensions across all ranges of effect sizes. We also compared the results of the shuffling method to a multivariate analysis of variance (MATLAB manova1). The permutation method and the MANOVA resulted in a similar estimation of the dimension. We preferred the permutation method since it can be calculated with less trials and does not assume data are drawn from a multivariate normal distribution.

### Effect Size

To calculate the coding of a variable throughout the entire epoch, we calculated the partial omega square (ω_p_^2^)^41–43^ effect size. The partial effect size measures how much of the variability of a neuron is related to the experimental variable in comparison to the variability that cannot be explained by any of the variables. Here we used ω^2^ to control the analysis of dimensions for the difference we recently found between populations in the sizes of neuron responses ^19^. Specifically, we calculated the number of spikes in 100 ms bins, 0 to 1200 ms after an event during the trial (target motion). We fitted an ANOVA model that included the direction of target motion as a variable, with the addition of time (the specific bin the sample came from). We used the following formula:

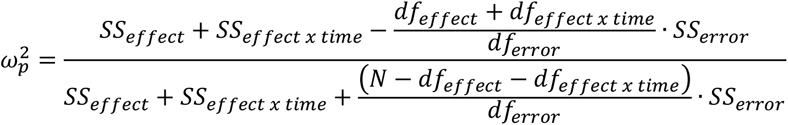

where *SS_effect_* is the ANOVA sum of squares for the effect of a specific variable, *SS_error_* is the sum of squares of the errors after accounting for all experimental variables, *df_effect_* and *df_error_* are the degrees of freedom for the variable and the error respectively and *N* is the number of observations (number of trials x number of time bins). *SS_effect x time_* is the ANOVA type II sum of squares for the interaction of a specific variable (i.e., target direction) with time, and *df_effect x time_* are the corresponding degrees of freedom. We included the interaction term since it quantifies the time-varying coding of the variable.

## Appendix

Let *X* be the activity of a neuron where the activity is composed of a signal and a noise term:

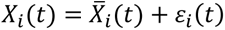

where *t* is the time on a trial, X(*t*) is the average activity in a task condition *i* and corresponds to the signal term, and *ε_i_*(*t*) is the noise term. We assume the following assumptions regarding the distribution of the noise term:

1. Zero average; i.e., 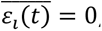, for all *t*.
2. Is uncorrelated with the signal across time; i.e., *COV*(*X_i_*(*t*), *ε_i_*(t)) = 0.
3. Noise terms across different task conditions are uncorrelated; i.e., *COV*(*ε_i_*(*t*), *ε_j_*(*t*)) = 0.
4. Noise is drawn independently across trials.

The calculations below are always performed over time, so for brevity, we omit the *t* in the equations.

Since noise and signal are uncorrelated and noise is drawn independently across trials the covariance between activity in two trials from two task conditions is equal to the covariance of the average activity:

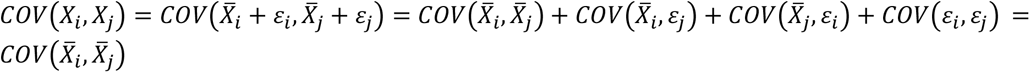

The correlation coefficient between two trials is defined as:

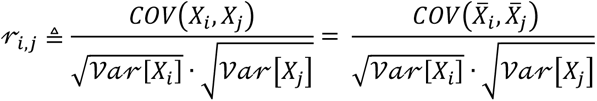

where *i* and *j* correspond to two task conditions.

The *Between* score is the correlation coefficient for trials in which *i ≠ j*.

When calculating *r*_ij_ for different trials in the same condition (*i* = *j*) it can be written as:

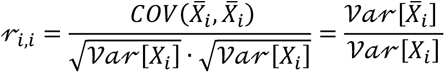

The *Within* score is defined as:

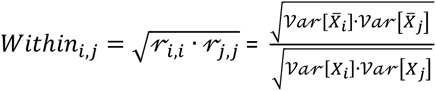

Therefore, the ratio between scores is:

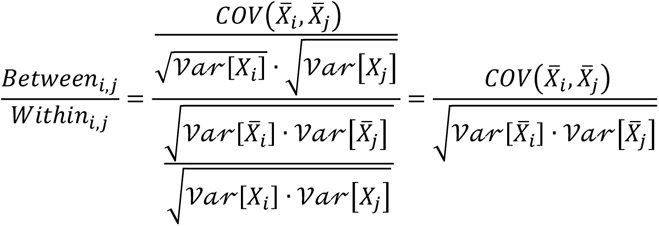

This means that the ratio of the *Between* to *Within* scores is equal to the correlation coefficient between the signal (average response) of the activity of the two neurons.

For each pair of trial conditions, we estimated the within and between scores by averaging their values across all possible pairs of trials, excluding the correlation of a trial with itself.

## Acknowledgments

This work is dedicated to the memory of Mrs. Lily Safra, a great supporter of brain research. This project received funding from the European Research Council (ERC) under the European Union’s Horizon 2020 research and innovation program (grant agreement No 755745) and the Israel Science Foundation (380/17).

---

**Figure S1:**
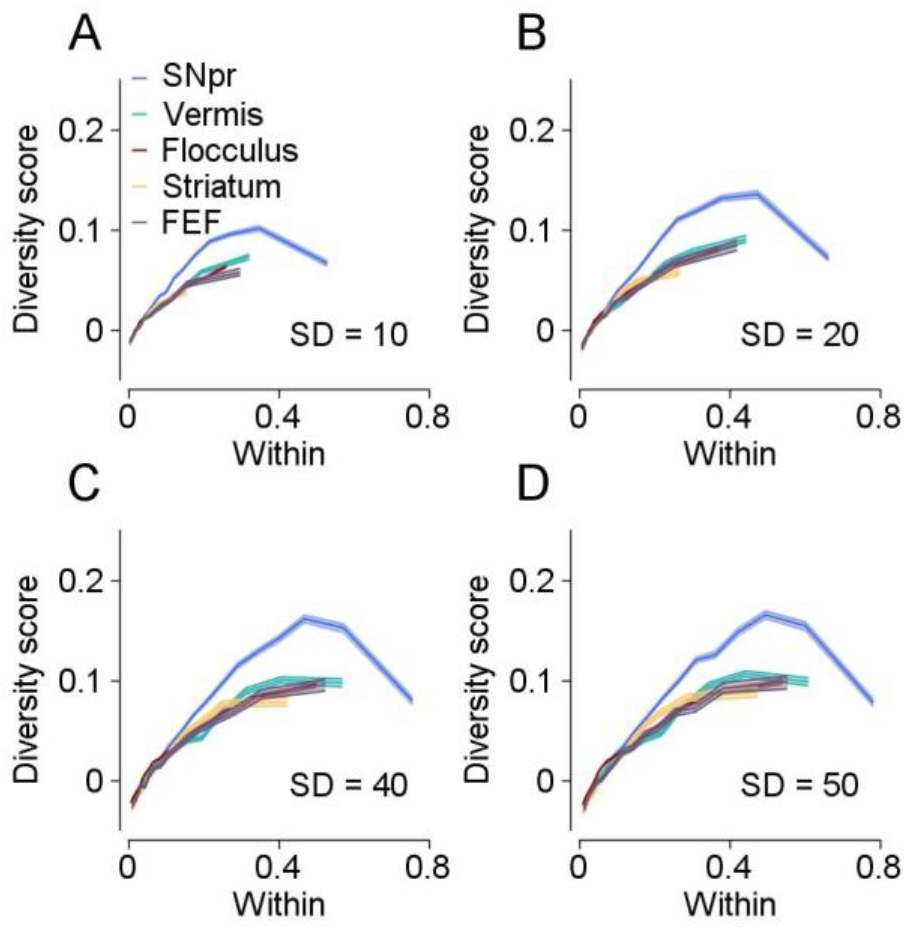
Diversity score results for different smoothing windows. **A-D** Within (horizontal axis) versus the Diversity score (vertical axis) for all populations in a range of standard deviation (SD) values of the Gaussian filter. The color indicates the population of neurons.

**Figure S2:**
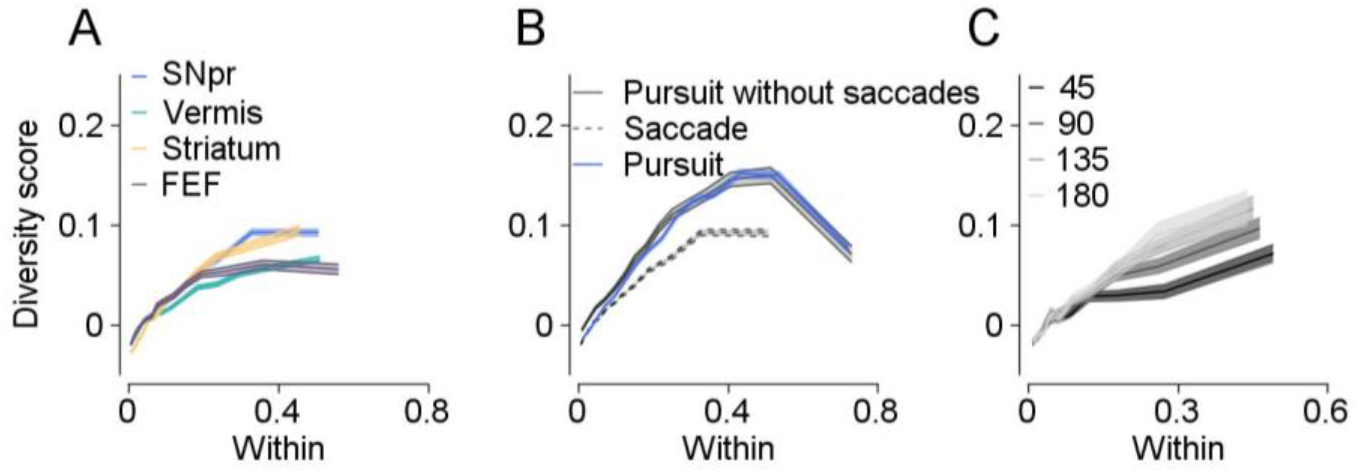
Diversity score for saccades. **A.** Same as Figure 3A for the saccade task. Within (horizontal axis) versus the Diversity score (vertical axis). Colors indicate the population of neurons (SNpr vs. Striatum p=1.1e-16, SNpr vs. Vermis p=5.7e-71, SNpr vs. FEF p=1.6e-04 Wilcoxon signed-rank test, see Methods). **B.** Control for catchup saccade during pursuit. SNpr during pursuit (blue) and saccade (black dashed line) and pursuit after removing catch-up saccades (black solid line). The time of the saccades was removed and treated as missing data. **C.** Same as Fig. 2C for saccade task. Comparison of the distances in target motion angle. The color of the trace shows the angle between the directions of eye movement.

**Figure S3:**
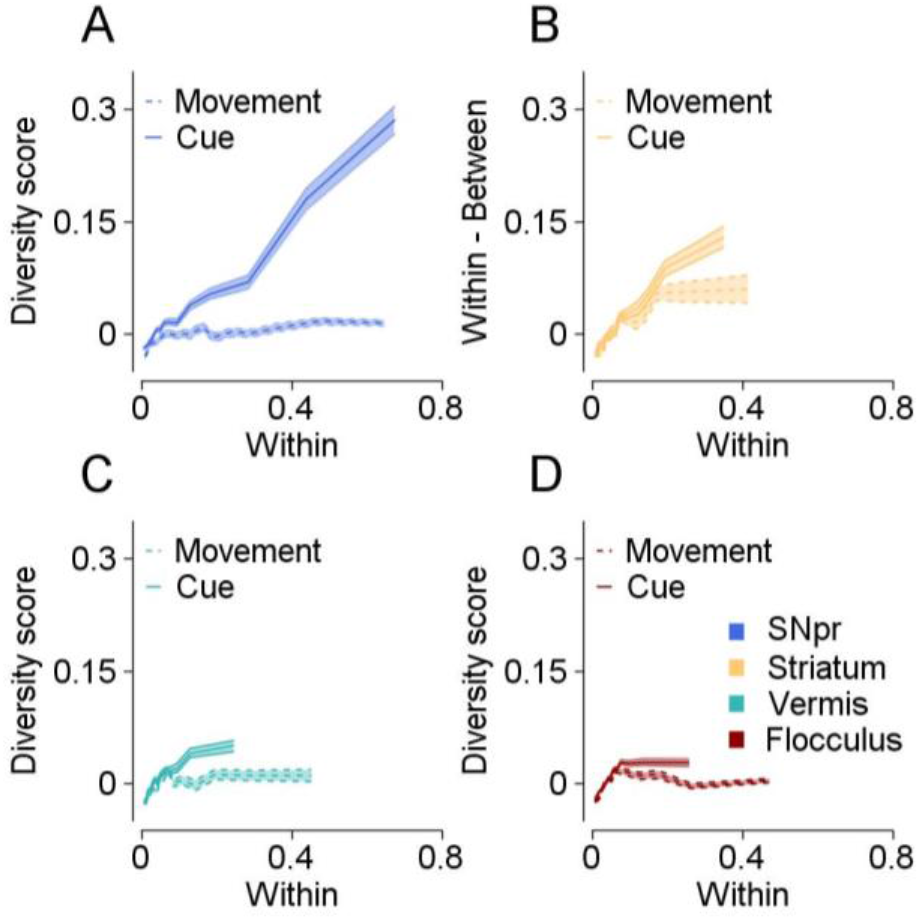
Temporal diversity of responses to the reward probability during cue and movement for different populations. Within (horizontal axis) versus the Diversity score (vertical axis). Colors indicate the population of neurons (blue- SNpr, yellow- Striatum, green- Vermis, red-Flocculus). Solid and dashed line show analyses for cue and movement.

**Figure S4:**
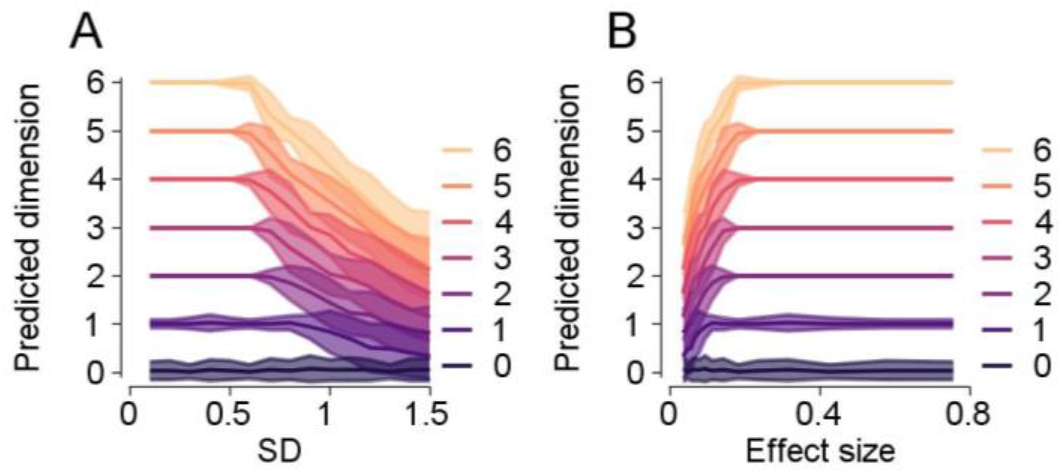
Simulation for validating the dimension analysis. **A.** The predicted dimension (vertical axis) as a function of the standard deviation of the noise (horizontal axis). The color indicates the true dimension, lines show averages, and bands the standard deviation across simulations. **B.** The predicted dimension (vertical axis) as a function of the effect size of the neuron (horizontal axis). Colors are same as in **A**.

**Figure S5:**
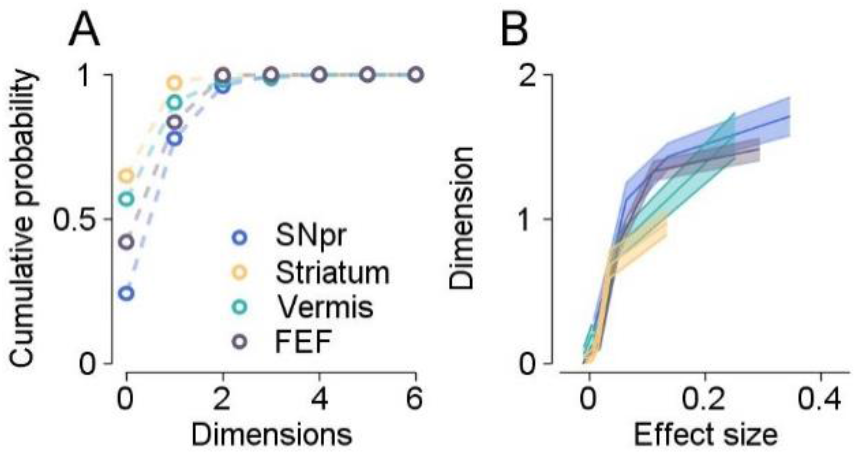
Dimensionality of the SNpr and other populations during the saccade task. **A.** Cumulative distribution of the dimension of the neurons. Colors indicate different populations (SNpr vs. Striatum p=5.4e-13, SNpr vs. Vermis p=1.3e-7, SNpr vs. FEF p=1.5e-3, Wilcoxon rank-sum test). **B**. Dimension of neurons (vertical axis) as a function of effect size (horizontal axis). Colors same as **A** (SNpr vs. Striatum p=9.0e-3, SNpr vs. Vermis p=0.74, SNpr vs. FEF p=2.9e-3 Wilcoxon signed-rank test, see Methods).

**Figure S6:**
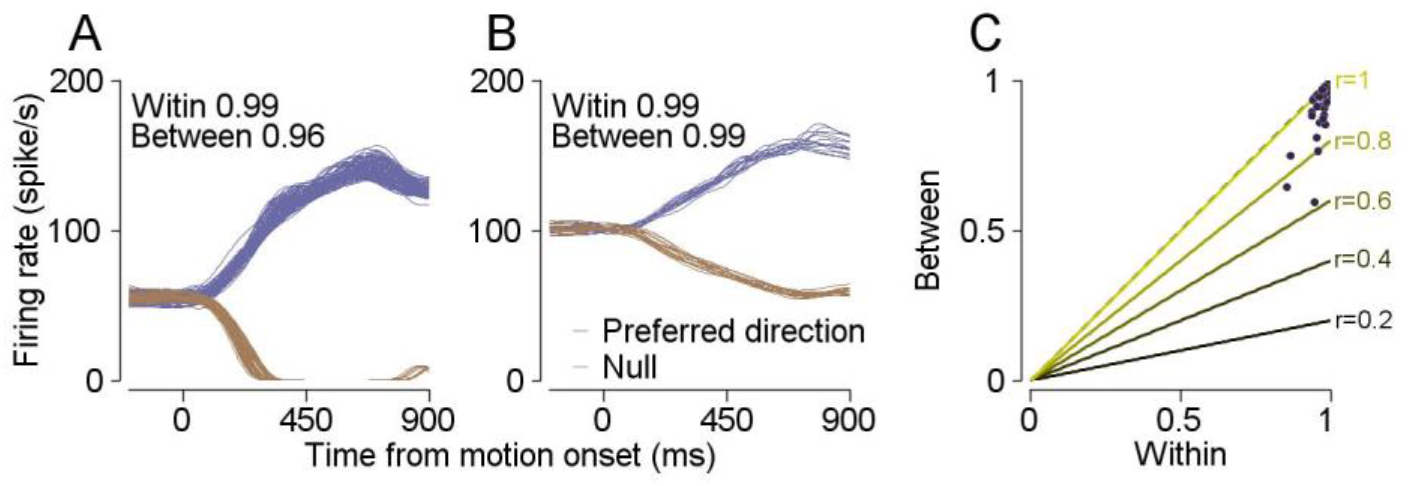
Diversity of motor neurons during pursuit. **A,B.** The firing rate (vertical axis) of two example motoneurons during pursuit eye movement in the preferred (purple) and null (brown) directions. The horizontal axis is the time from motion onset. Each thin trace shows the smoothed firing rate on a single trial (SD = 30 ms). The responses of motoneurons in the null and preferred direction tended to be highly correlated. Truncation at zero resulted in some non-linearity between the null and preferred directions. **C.** Within vs. Between scores for motoneurons. Each dot shows the scores calculated across the preferred and null directions of a single neuron. Solid lines indicate different theoretical correlation slopes. Analysis conducted on 49 abducens motoneurons (36 and 13 from monkeys P and I) used in previous studies^44, 45^.

**Figure S7:**
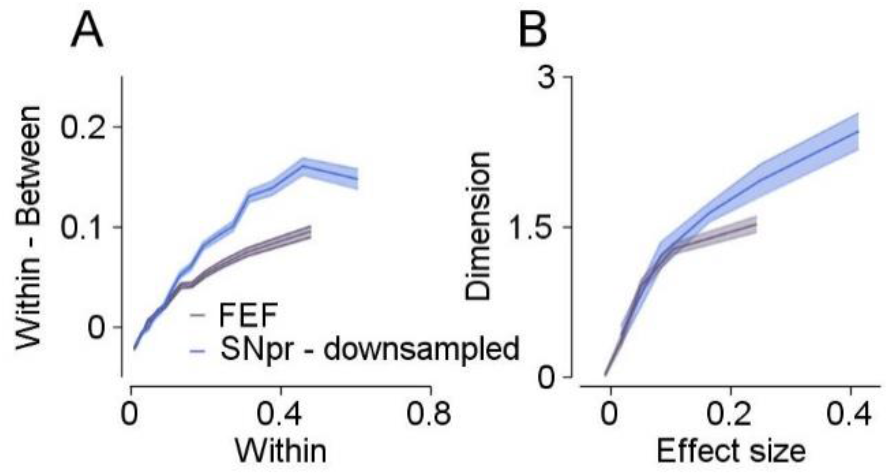
Matching the trial numbers for Within-Between and dimension analysis. To control for the different numbers of trials in the FEF population, we down-sampled the number of trials for each direction in the SNpr population to match the FEF population. Colors indicate the population of neurons. **A.** Within (horizontal axis) versus the Diversity score (vertical axis) (SNpr vs. FEF p=2. 2e-24, Wilcoxon rank-sum test). **B.** Dimensionality as a function of the effect size (SNpr vs. FEF p=1.2,e-4 Wilcoxon rank-sum test).

